# Performing highly parallelized and reproducible GWAS analysis on biobank-scale data

**DOI:** 10.1101/2023.08.08.552417

**Authors:** Sebastian Schönherr, Johanna Schachtl-Riess, Silvia Di Maio, Michele Filosi, Marvin Mark, Claudia Lamina, Christian Fuchsberger, Florian Kronenberg, Lukas Forer

## Abstract

**Motivation:** Genome-wide association studies (GWAS) in large biobanks are transforming genetic research and enable the detection of novel genotype-phenotype relationships. In the last two decades, over 60,000 genetic associations across thousands of human diseases and traits have been discovered using a GWAS approach. Due to denser genotyping and increasing sample sizes, researchers are increasingly faced with computational challenges when executing GWAS analysis. A reproducible, modular and extensible pipeline with a focus on parallelization is essential to simplify data analysis and to allow researchers to devote their time to other essential tasks such as result interpretation and downstream analysis.

**Results:** Here we present nf-gwas, a Nextflow pipeline to run biobank-scale GWAS analysis. The pipeline automatically performs numerous pre- and post-processing steps, integrates regression modeling from the REGENIE package and currently supports single-variant, gene-based and interaction testing. nf-gwas also includes an extensive reporting functionality that allows to inspect thousands of phenotypes and navigate interactive Manhattan plots directly in the web browser. The pipeline is extensively tested using the unit-style testing framework nf-test to ensure code maintainability, a crucial requirement in clinical and pharmaceutical settings. Furthermore, we validated the pipeline against published GWAS datasets and benchmarked the pipeline on high-performance computing and cloud infrastructures to provide cost estimations to end users.

**Availability:** nf-gwas is free available at https://github.com/genepi/nf-gwas.

## Introduction

Over the last decades, genome-wide association studies (GWAS) have emerged as a key technology to discover insights into the genetic architecture of diseases. The GWAS approach shows remarkable success and identified so far more than 60,000 genetic associations ^1^. Nevertheless, increasing sample sizes and phenome-wide association studies (PheWAS), with thousands of phenotypes analyzed for one specific genomic variant, require a continuous improvement of methods to run GWAS regression models in a computationally efficient way. Over the last few years, novel and faster machine learning methods especially for the PheWAS use-case have been published, which are ideally suited for high parallelization ^2^. While the availability of new GWAS methods is critical for the success of large-scale analyses, running a GWAS analysis on a computational infrastructure still involves many technical and workflow-specific challenges. First, GWAS analysis consists of several necessary pre- and post-processing steps to improve result quality. This ranges from simple file-format checks to time-intensive file conversions between different genetic formats. For example, the REGENIE software currently does not accept input datasets in VCF format, the default export format from popular imputation servers ^3^. Therefore, researchers are forced to invest their time in script preparation for data manipulation. Second, while novel methods such as REGENIE allow an efficient parallelization on e.g., a chromosome level using CPU threads, researchers have to execute a higher parallelization (e.g. by genomic chunks) manually. This requires detailed documentation of executed steps to guarantee the reproducibility of results and is especially challenging when working in a distributed environment such as local high-performance computing (HPC) or cloud environments where data needs to be distributed to individual compute nodes. Third, generated GWAS datasets can be large and require the visualization of millions of data points or comparison of phenotypes in a user-friendly and scalable way. Due to all these reasons, researchers often have to dedicate a substantial part of their time to technical and workflow issues instead of focusing on the biological interpretation of results. Furthermore, these manual processes and individualized solutions are error-prone and do not promote reproducibility of results. To overcome these problems, pipelines are a prominent way to execute GWAS ^4–6^. While these pipelines are a step forward, they do not include features for biobank-scale datasets, different association testing modes, post-processing steps (e.g. lift-over) or scalable visualizations. Furthermore, workflow code is often untested, which complicates its usage and extensibility in critical settings.

Here we present a versatile computational pipeline to perform highly scalable and reproducible GWAS analysis in distributed environments. The pipeline is based on the popular Nextflow workflow manager ^7^ and supports researchers to run GWAS on thousands of phenotypes in parallel. The pipeline includes all necessary pre- and post-processing steps, currently builds on the REGENIE method for running the regression models and has been developed with a focus on high parallelization including thousands of samples and phenotypes. The pipeline also supports various association testing modes such as classical single-variant but also gene-based and interaction testing. This makes it suitable for diverse scientific questions that will likely become more important in the future, as not only imputed SNPs but also whole exome sequencing (WES) and whole genome sequencing (WGS) data become available in more and larger cohorts ^8, 9^. To validate our pipeline, we reproduced published GWAS results showing excellent agreement between approaches. The pipeline has also been extensively used within our institute for over 2 years ^10^, has been improved by many contributions from the open-source community and uses the popular nf-test testing framework for unit-style testing (see Resources). To provide users benchmarks and cost estimations, we run the pipeline in widely used environments (HPC Slurm cluster, AWS Batch Cloud) using different parallelization strategies and chunking levels. Overall, nf-gwas is a best-practice and well-tested workflow for performing GWAS analysis and can be run with the power of Nextflow in any distributed environment.

## Methods

### Design and Implementation

The nf-gwas pipeline is implemented as a Nextflow ^7^ DSL2 pipeline. The implemented processes are massively parallelized on chromosomes or genomic chunks with the overall goal to provide an efficient utilization of computational resources. All dependencies used by the pipeline are available in Docker or Singularity containers providing transportability. The modular architecture of nf-gwas enables an easy integration of future updates and extensions to support emerging GWAS methodologies. nf-gwas provides several Nextflow profiles allowing users to execute GWAS on various computing infrastructures. Moreover, we used nf-test to ensure that the pipeline code works correctly with no errors, which also improves its robustness and maintainability against updates.

### Parallelization

nf-gwas allows a high parallelization of both REGENIE steps (see Figure 1). Step 1, which fits the whole genome regression model to the traits, has been parallelized using the best-practices approach provided by the authors of REGENIE. First, the level 0 models are run on a chunk-level and are then merged in parallel by phenotype. Step 2, which consists of the regression model, runs by default on a chromosome level. For HPC or cloud computing, splitting chromosomes into smaller chunks can be beneficial. This allows lowering the per-chunk memory usage and enables a better distribution across a large number of computational nodes. To flexibly adjust the type of parallelization, nf-gwas provides, besides the chunk size parameter, also the possibility to specify the applied chunk strategy. Chromosomes can be split either by a genomic range (default) or by specifying the number of variants for each chunk. After the whole genome regression model is applied, the pipeline operates on a phenotype level for better runtime estimations. We evaluated the different chunk strategies and chunk size values in the Results section.

**Figure 1:**
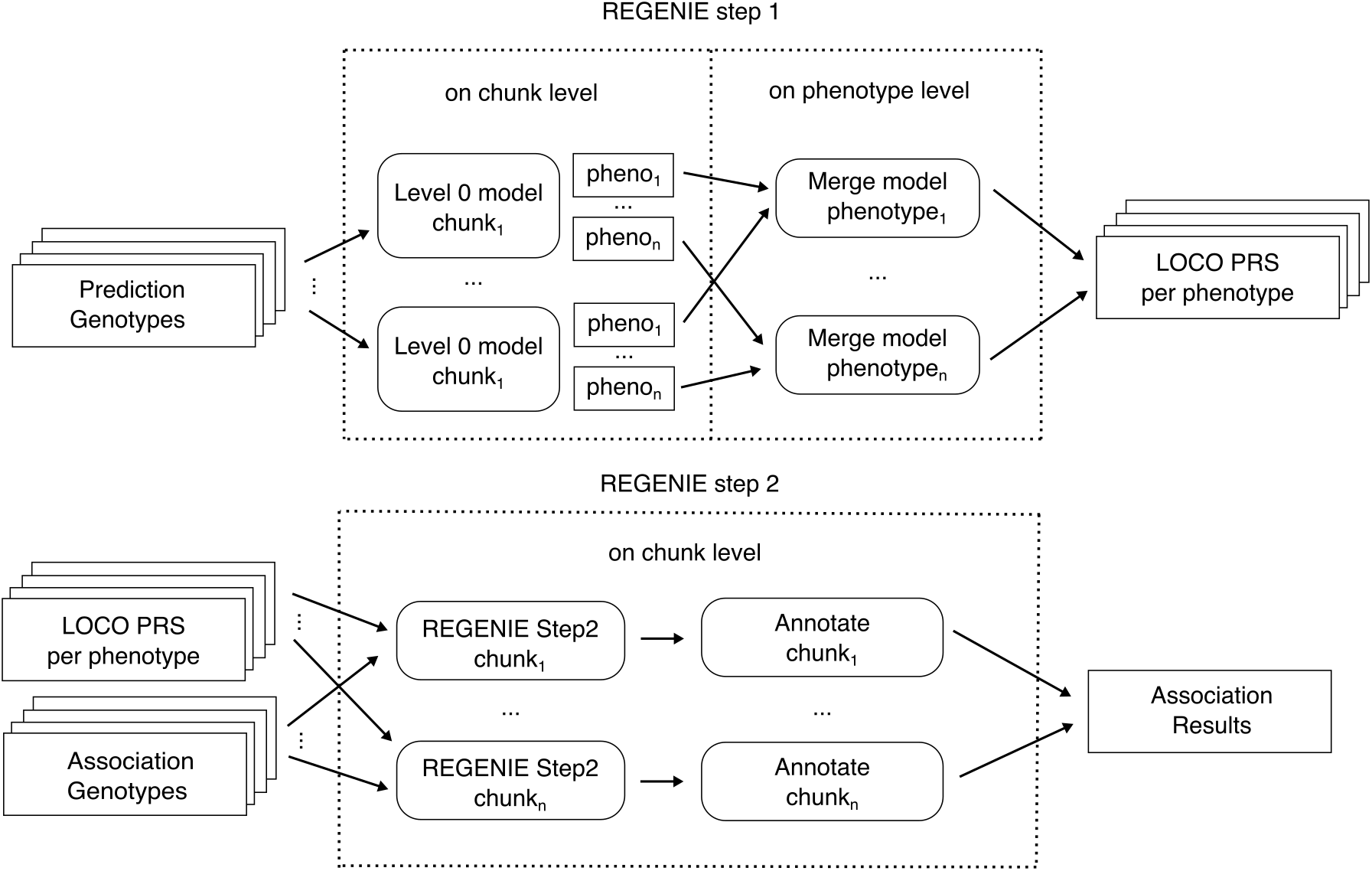
Overview of the REGENIE parallelization. Prediction genotypes (Step 1 Level 0) are first run on a user-defined chunk level and calculated predictions are then merged by phenotype in parallel. Step 2 (association of genotypes with phenotypes) and annotation is performed in parallel for all association genotype chunks in combination with the LOCO predictions. Afterwards, annotated association results are merged per phenotype and all post-processing steps within the pipeline are performed in parallel for all phenotypes (not shown).

### Annotation and Interactive Visualizations

Summary statistics are annotated with nearest genes and rsIDs using our genomic-utils library in parallel for all genomic chunks (see Resources). The library uses an annotation strategy similar to available tools (e.g. LocusZoom, PheWeb). To prepare the required reference files, we downloaded the gene annotations (hg19 and hg38) from GENCODE release v32 ^11^ and filtered them for common gene types (protein_coding, IG_C_gene, IG_D_gene, IG_J_gene, IG_V_gene, TR_C_gene, TR_D_gene, TR_J_gene, TR_V_gene; for details on gene/transcript biotypes). Variants are annotated with the nearest gene, its start and end position and the distance to the respective gene. If a variant is located within two or more overlapping genes, all genes are reported. For annotation with rsIDs, a reference file based on dbSNP v154 was created and an index was built to enable a fast lookup. To avoid numerous downloads of the file, a local tabix-indexed reference file can be specified in the Nextflow configuration.

Both an interactive and static report are generated in parallel for each phenotype. All reports are collected in an HTML overview report which simplifies navigation between phenotypes. The interactive Manhattan plot is created based on a binning algorithm similar to LocusZoom ^12^ that supports millions of variants. By default, top loci are defined +/-200 kb around the most significant SNPs that reach by default at least a genome-wide significance of -log_10_(p)>-log_10_(5*10^-8^). The detailed report is generated using R v4.1.0 and RMarkdown.

### Validation

REGENIE has already been successfully validated against other GWAS software elsewhere ^2^. However, to ensure that our parallelized implementation works correctly, we validated our pipeline with UK Biobank data against (a) publicly available summary statistics for apolipoprotein B (apoB; data field: 30640) and total cholesterol (data field: 30690) for European ancestry by Pan-UK Biobank and (b) selected SNPs per locus associated with lipoprotein(a) [Lp(a)] (data field: 30790) from a GWAS by Said et al. ^13^.

We downloaded available summary statistics from Pan-UK Biobank and compared p-values using Miami plots. Briefly, the downloaded summary statistics were generated using SAIGE implemented in Hail Batch on rank inverse normally transformed (RINTed) phenotypes, using ancestry defined based on genetic data (European n=399,003 for apoB and n=400,963 for cholesterol). The Pan-UK Biobank GWAS was performed for 23,861,713 SNPs for apoB and 23,861,710 SNPs for cholesterol, adjusted for age, sex, age*sex, age^2^, age^2^*sex and the first 10 principal components (PCs). Our GWAS was based on nf-gwas v1.0.0 using REGENIE on RINTed phenotypes, European ancestry was defined by data field 21000 (European n=375,278 for apoB and n=377,097 for cholesterol) and GWAS was adjusted for age, age^2^, sex, genotyping batch and the first 30 PCs. For step 1 of REGENIE we used directly genotyped variants that were filtered for a minor allele frequency (MAF) ≥ 0.01, minor allele count (MAC) ≥ 100, genotype missingness<0.1, Hardy-Weinberg equilibrium test p ≥ 10^-15^ and sample missingness < 0.1 within the pipeline. For step 2 of REGENIE we included variants imputed to a custom reference-panel (hg19) with MAC > 100 and an imputation info score > 0.3 (apoB 33,390,477 and cholesterol 33,439,525 SNPs).

For validation against the Lp(a) GWAS by Said et al., we compared effect estimates and p-values using correlation and Bland Altman plots and calculated the correlation of selected SNPs since full summary statistics are not publicly available. Data were available from Said et al. Supplemental Table II ^13^. Briefly, Said et al. performed GWAS (19,000,000 SNPs) in all ancestries in UK Biobank (n= 371,212) using BOLT-LMM v2.3.1, RINTed serum Lp(a) concentrations and the analysis was adjusted for age, age^2^, genotyping array, lipid-lowering drug usage and the first 30 PCs. We performed the GWAS based on nf-gwas v1.0.0 using REGENIE on RINTed Lp(a) including all ancestries (n=371,458) and GWAS was adjusted for age, age^2^, genotyping batch, statin treatment and the first 30 PCs. For step 1 of REGENIE we used directly genotyped variants that were filtered for a MAF ≥ 0.01, MAC ≥ 100, genotype missingness < 0.1, Hardy-Weinberg equilibrium test p ≥ 10^-15^ and sample missingness < 0.1 within the pipeline. For step 2 of REGENIE we included variants imputed to a custom reference-panel (hg19) with MAC > 100 and an imputation info score > 0.3 (42,188,331 SNPs).

### Evaluation and Compute Infrastructure

Experiments were performed on two different computing environments. All validation experiments and benchmarks for different chunk sizes and chunking strategies, were executed on an in-house HPC cluster. The in-house HPC cluster is based on Slurm with 60 CPU nodes (AMD EPYS 7713P) consisting of 64 cores and 1TB RAM each. Local experiments were executed with the appropriate Nextflow profile (--profile slurm) using Nextflow version 22.10.4 in combination with Singularity. To provide cloud cost estimations, we ran experiments on AWS Batch with 450 vCPUS and different instances types (on demand, spot) executed with nf-tower (see Resources). As a dataset we used data from the UK Biobank under application number 62905 with ∼460K samples and >90M variants and 100 copies of the Lp(a) phenotype to mimic running the pipeline on several phenotypes in parallel.

## Results

### Pipeline Overview

The nf-gwas pipeline automatically performs three main steps including (a) data pre-processing steps such as quality control, phenotype validations or file conversions, (b) alternative association analyses using REGENIE ^2^ (currently single-variant, interaction and gene-based mode) and (c) data post-processing steps such as annotation, interactive visualization, indexing and optional lift-over to allow efficient data exploration and further processing of results (Figure 2). Parameters for the mandatory and optional steps can be defined in a Nextflow configuration file. The pipeline supports typical file formats for association testing (VCF, bgen) and different genome builds (hg19/hg38) to avoid error-prone and time or resource intensive conversions. To reduce costs and to efficiently use computational resources, processes are massively parallelized (see Methods). Optionally, lift-over of results to other genome builds (hg19/hg38) is performed to e.g. allow out of the box meta-GWAS with studies imputed to other reference panels. An overview of all required and optional parameters can be found in our online documentation (see Resources). The documentation also provides instructions on executing the pipeline for advanced users (see “Getting started”) and for users without prior command line knowledge (see “Beginners guide”).

**Figure 2:**
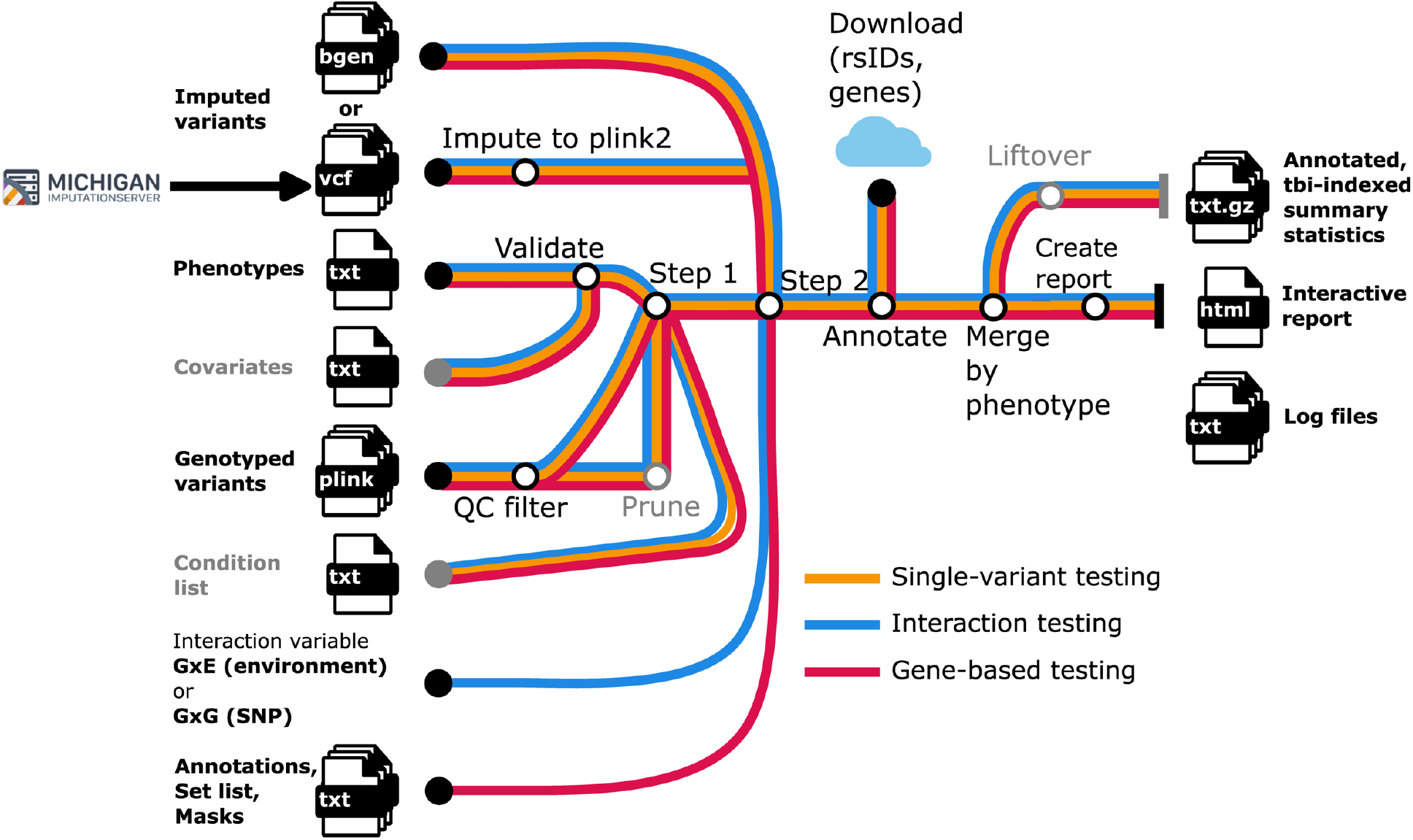
Overview of the nf-gwas pipeline. Input files are on the left side and output files on the right side. The color defines the type of mode. Optional steps are marked in gray.

### Pre-processing: Input Validation and Quality Control

In the pre-processing step, the file format of the phenotypes and the optional covariates is validated. If phenotypes are not normally distributed, nf-gwas provides a parameter to rank inverse normally transform (RINT) phenotypes and it displays the distribution of the RINTed phenotype in the final report. If the genetic data is provided in VCF format, it is converted to plink2 to be compliant with REGENIE input format. To account for relatedness and population structure, REGENIE first uses a subset of genetic markers in step 1 to fit a whole genome regression model that is then used as covariates according to the leave one chromosome out scheme in step 2 (i.e. tests the association of a larger set of genetic variants, e.g. imputed SNPs, with the phenotypes) ^2^. For step 1, REGENIE requires directly genotyped SNPs with a good quality that should optimally be around 500,000 SNPs and should not exceed 1,000,000 SNPs to avoid a high computational burden. Therefore, a set of filters are automatically applied using plink2 ^14^ with default values currently set to: MAF ≥ 0.01, MAC ≥ 100, genotype missingness < 0.1, Hardy-Weinberg equilibrium test p ≥ 10e-15 and sample missingness < 0.1 (adjustable in the Nextflow configuration file). Optionally, the variants can further be pruned using plink2 with adjustable thresholds and filters.

### Association testing

The pipeline currently supports three analysis modes including (a) single-variant (default), (b) gene-based and (c) interaction testing with genetic and environmental variables. All three modes can be run on thousands of phenotypes in parallel. Further, all three modes support adjustment for a set of covariates directly within the pipeline and conditioning on a set of genetic variants that are also added as covariates in step 1 and step 2 of REGENIE. The pipeline works for quantitative and binary traits and additive, as well as dominant and recessive association testing. Exemplary data input files can be found in the GitHub repository.

### Post-processing: Annotation, Reporting and Data Manipulation

The pipeline automatically performs post-processing of data including (a) annotation, (b) visualization of results in an interactive report and (c) data preparation for downstream analyses.

For annotation, results are automatically linked with rsIDs, the name of the nearest genes and the distance to those genes. Top gene loci are displayed with links to the GWAS catalog. For visualization of results, an interactive report is automatically generated that allows exploration of results. The overview report provides a summary on the number of top loci and allows the navigation of phenotype-specific reports (Figure 3 left side). For the selected phenotype, the overview report allows the navigation to “Interactive Plots” and “Details and Phenotype” (Figure 3 tabs above phenotype name). The tab “Interactive Plots” can be used to interactively explore the GWAS results. Hovering the mouse over a variant will depict more details in a blue box. The report also contains a table of the top loci with the most important information and links of (a) the lead SNP and (b) the annotated nearest gene to the GWAS catalog to quickly look up important details and published findings. The tab “Details and Phenotype” contains a static Manhattan plot and q-q plot suitable for publications, a more detailed top loci table (e.g. also including allele frequencies, imputation info score and distance to the nearest gene), summary statistics on the phenotype and project summary including pipeline version.

**Figure 3:**
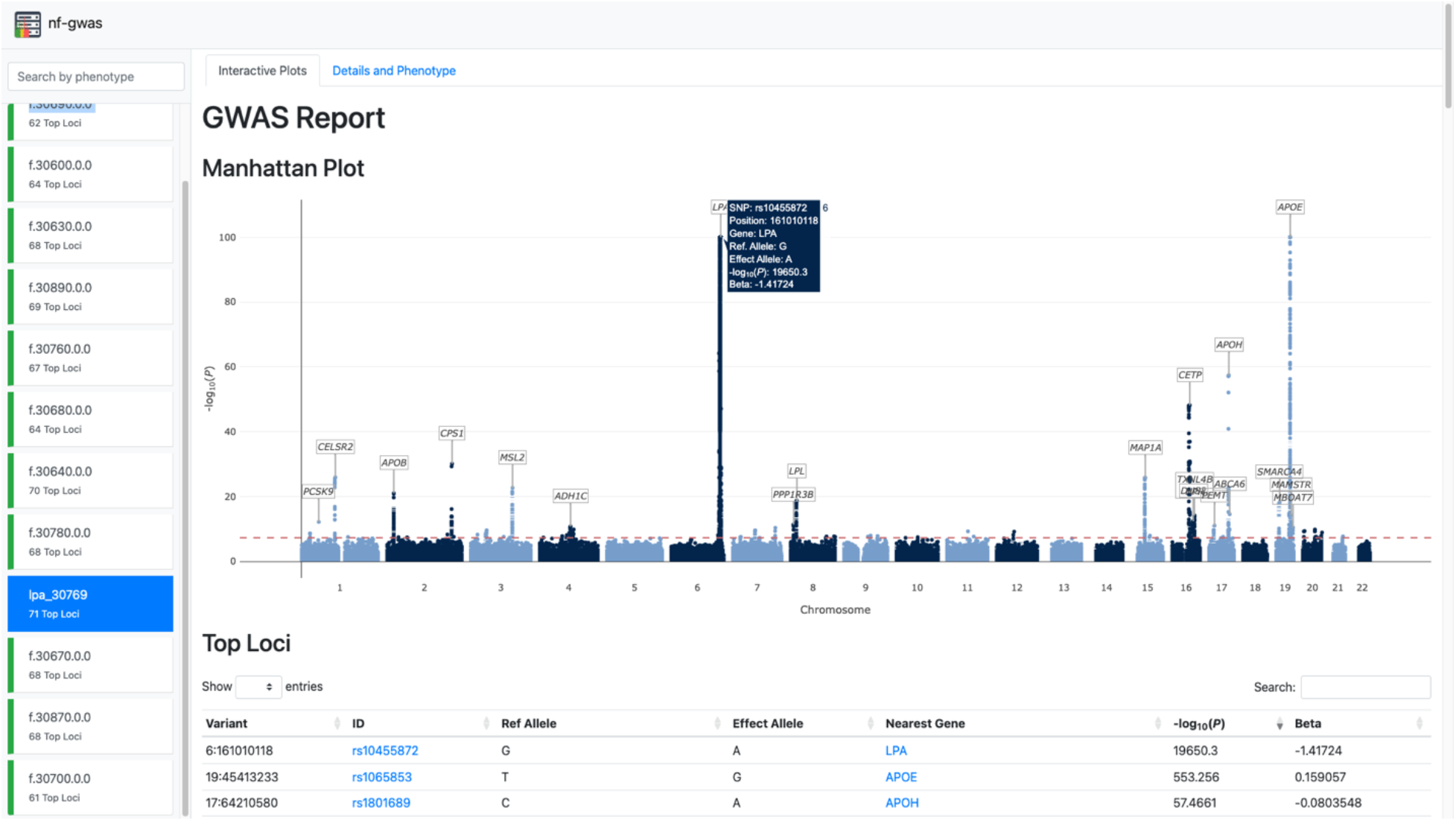
Example of a nf-gwas report. The index gives an overview on all phenotypes included into the run and the number of top loci found (left side). In the tabs at the top users can navigate between the interactive plots and more details on the respective phenotype (e.g. phenotype summary statistics, high resolution Manhattan and QQ plots, summary statistics on the phenotype and log files). By hovering over the interactive Manhattan plot, more details about the variants are depicted in a box (e.g. here for rs10455872). The top loci table below is searchable and rsIDs and gene names are linked to the GWAS catalog.

For downstream analyses, summary statistics are automatically tabix-indexed and are compatible with LocusZoom and LocalZoom ^12^. Optionally, the pipeline can perform automatic conversion of GWAS results into a target build, if required for further downstream analyses (e.g. meta-GWAS with studies on another genome build).

### Output files and structure

Our pipeline produces several output files. The overview HTML report (or index report) summarizes all main results (e.g. interactive Manhattan/q-q plot, phenotype statistics) from the different traits in a browsable way. Individual phenotype reports are provided (folder index_report) including publication-ready plots. Summary statistics are included in the results folder and consist of an unfiltered REGENIE file and a p-value filtered file (folder tophits) for each phenotype. Importantly, all pipeline executions are logged for reproducibility and log information and the validated input file are also provided within an output folder.

### Validation of GWAS results

We validated and benchmarked GWAS results of our parallelized pipeline against published data generated using (a) a pipeline with SAIGE implemented in Hail Batch by the Pan-UK Biobank project and (b) BOLT-LMM v2.3.1 applied by Said et al. ^13^. For the Pan-UK Biobank dataset, Miami plots show an excellent agreement between the publicly available summary statistics and the output of our pipeline (Figure 4). A complete comparison with Said et al. was not possible based on Miami plots since statistics were only publicly available for selected SNPs of the significant loci. Of the 39 selected SNPs, 37 were available in our summary statistics (Supplement Table S1). Correlation is high between both beta estimates (rho = 0.99, p<2.2*10^-15^) and -log_10_(p) values (rho 0.94, p<2.2*10^-15^; Supplement Figure S1 A, C, E). Bland Altman plots also show a good agreement between the GWAS of Said et al. and our pipeline (Supplement Figure 1 B, D, E) except for slightly larger differences for *LPA* (Said et al. versus nf-gwas: beta = -1.02 versus -1.42 and -log_10_(p) = 19379 versus 19594, respectively; Supplement Table S1). Both experiments show that our pipeline gives out highly reliable results even across GWAS models independently of the applied chunking strategy, which resulted in a much faster wall-time compared to chromosome-wise execution.

**Figure 4:**
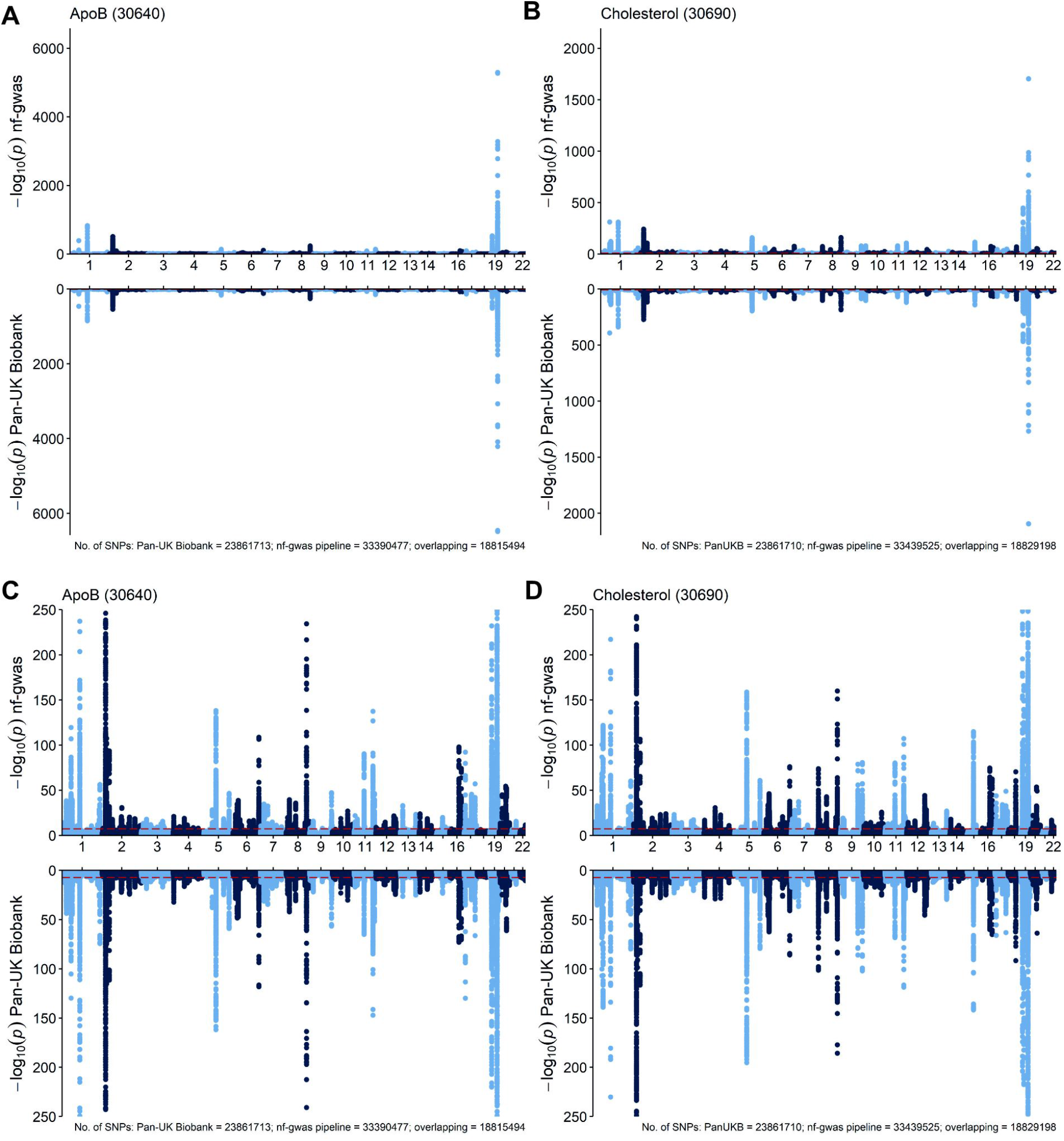
Miami plots of benchmarking to Pan-UK Biobank summary statistics. The red dashed line marks genome-wide significance. Plots **C** and **D** are cut at -log_10_(p) = 250 excluding SNPs with higher values.

### Parallelization Benchmark

We performed a GWAS analysis using the UK Biobank imputed dataset (90M input variants, 10 phenotypes) on our local HPC infrastructure (see Methods). We executed 7 experiments consisting of different chunk strategies and chunk sizes. Each experiment has been executed 3 times, whereas the mean time is reported in Table 1. We show how the different parallelization strategies affect the overall runtime. While a chromsome-based execution can be achieved easily by user-specific scripts, the chunk-based execution and therefore high parallelization has been implemented directly within nf-gwas and results in better utilization of large cluster architectures. Our chunksize experiments show that for both strategies (variants, range) a value which results in 567 chunks (chunksize of 5M for range, chunksize of 168,200 for variants) results in the best overall performance (4h 35 min for the variants strategy, 4h 48 min for the range strategy). Our results further show, that an execution using the variants strategy results in a more uniform execution time for all 567 chunks (median/maximum: 28 min / 36 min) compared to the range strategy (median/maximum: 28 min / 58 min).

**Table 1:** GWAS Runtime and CPU hours on a local HPC Slurm cluster. We executed a GWAS using a UK Biobank consisting of ∼460K samples, > 90M variants and 10 quantitative phenotypes. Each experiment consists of a different chunking strategy and chunk size. Each experiment was performed three times and the mean is reported.

| Chunks | Chunksize | Type | Local HPC |  |
| --- | --- | --- | --- | --- |
|  |  |  | Runtime | CPU hours |
| 5432 | 500K | region | 5h 56 min | 2,196 |
| 5432 | 17,170 | variant | 5h 27 min | 1,491 |
| 567 | 5M | region | 4h 48 min | 2,163 |
| 567 | 168,200 | variant | 4h 35 min | 2,195 |
| 293 | 10M | region | 5h 14 min | 2,184 |
| 293 | 330,500 | variant | 5h 19 min | 2,180 |

### Portability to AWS Batch

To show the biobank-scale possibilities of nf-gwas, we performed a full GWAS analysis using the UK Biobank imputed dataset (460,000 samples, 90M input variants, 3TB of data) with 100 phenotypes with AWS Batch. Overall, nf-gwas performed the GWAS analyses in 15 hours using 4,600 CPU hours with estimated costs of ∼$130 using AWS spot instances. This corresponds to a cost of 1 US Dollar for more than 3,000 samples running on 100 phenotypes in parallel and includes all pre- and post-processing steps. We further investigated the effect of using on-demand versus spot instances. While AWS on-demand instances have a fixed price per hour and up-time is therefore assured, spot instances are cheaper but can be interrupted depending on the current availability of the instances. To support spot instances, we implemented an interruption strategy to avoid that an analysis fails in case a single spot instance is interrupted. Table 2 shows the price of a GWAS analysis running on 5.76M variants (chromosome 6), ∼460,000 samples and 100 phenotypes. It can be seen that by using spot-instances ∼80% of overall costs can be saved. Overall, usage of spot instances can result in a longer overall runtime, but with the advantage of reduced costs.

**Table 2:** GWAS runtime and costs on AWS Batch. We executed a UK Biobank GWAS on ∼460K samples, 5.76M variants and 100 quantitative phenotypes. We executed the experiment using on demand and spot instances and show that up to 80% of the costs can be saved when using spot instances. We executed the experiments three times, since spot instance prices depend on available resources.

| Variants | Max. vCPUs | Instance Types | Costs (US Dollar) | CPUS hours | Runtime | Saving |
| --- | --- | --- | --- | --- | --- | --- |
| 5.76M | 450 | on demand | 34.331 | 553 | 2h 43 min | - |
| 5.76M | 450 | spot | 7.5 | 311 | 1h 59 min | 78 % |
| 5.76M | 450 | spot | 7.4 | 308 | 1 h 55 min | 78 % |
| 5.76M | 450 | spot | 7.6 | 311 | 1h 58 min | 78% |

### Portability to the UKB Research Analysis Platform (RAP)

The REGENIE method is also available on RAP as an application (within the tool library), which allows users to specify inputs and output parameters graphically. The application only includes the GWAS method itself and does not include any pre- or post-processing steps, which are available in nf-gwas. Since RAP does not allow to execute Nextflow workflows natively, users can still benefit from the nf-gwas pipeline on RAP by starting a single cloud workstation, downloading the data into the cloud workstation and running the pipeline. This setup will allow to use our well-test GWAS workflow without downloading any data.

## Discussion

While biobank scale datasets with millions of samples enable the discovery of novel genetic associations for diseases and traits, researchers are faced with new bioinformatic challenges at the same time. In the field of GWAS analysis, new methods such as REGENIE have been developed over the last years, offering memory efficient solutions for parallel execution of thousands of phenotypes ^2^. Still, the execution over multiple machines involves detailed and field-specific knowledge on distributed environments and can take a considerable amount of time. Different computational pipelines exist to simplify the execution over machines, nowadays often based on the Nextflow workflow system (Supplement Table S2) ^4–6^. These pipelines encapsulate necessary parallelization steps and provide users with a ready-to-use pipeline with the advantages of reproducibility and portability to other systems. Nevertheless, no pipeline currently exists solving the use cases many institutes like ours face. These include the highest possible parallelization of all data-intensive steps, visualization of millions of data points (e.g. Manhattan plot) directly in the browser, out-of-the-box support for different genetic formats (e.g. coming from Michigan Imputation Server and UK Biobank), or post-processing steps (e.g. lift-over) for usage in meta-GWAS analysis. Furthermore, no benchmarks for different environments or cost estimations for public clouds are currently available on how the pipelines perform e.g. on different parallelization levels.

Here we present nf-gwas, which includes a rich feature set and can be used for different use cases (such as single variant association testing, gene-based testing for rare variants or interaction-testing). Since rare variants are available in an increasing number of populations and large datasets also reach the statistical power to investigate interactions, gene-based testing and interaction analysis may become more relevant in the future and have already proven as important tools for novel findings ^8, 9^. Our pipeline is based on Nextflow and benefits from all its advantages ^7^. First, collaborators can use the pipeline locally to calculate summary statistics with a clearly defined workflow excluding the possibility to introduce artifacts or errors. Second, reproducibility of results is given at any time and collaborators can re-run experiments at a later point using the same software stack. Third, users get a clean and standardized project structure allowing them to navigate quickly between projects. Fourth, its modular structure allows it to add new GWAS methods in the future or execute optional steps for specific scenarios. For example, new imputation reference panels are built on different builds (currently hg19 or hg38) and an optional lift-over step allows the migration between builds. Finally, the pipeline also benefits from high quality code due to inclusion of unit-style testing provided by nf-test. This motivates collaborators to add new functionality without breaking existing ones. Our pipeline has been extensively validated and benchmarked. Validation ensures that our parallelized pipeline does not introduce artefacts or errors compared to published and already validated non-parallelized GWAS models. The benchmarks on AWS provides cost estimates on typical biobank-scale datasets to users. We show that a large GWAS including 460,000 samples, >90M variants and 100 phenotypes can be run in ∼15 hours, allowing to analyze more than 3,000 samples for 1 US Dollar.

The pipeline currently has some minor limitations. While REGENIE is currently the computationally most efficient method, it is also the only supported one in nf-gwas. Nevertheless, our modular structure will allow us or others to integrate new methods (e.g. fastGWA-GLMM ^15^) as a new process without re-writing the overall workflow. Future developments of the pipeline include forming a community behind nf-gwas e.g. by discussing the possibility to transform it into an nf-core pipeline or improving the functionality by adding additional steps for pre- or post-processing. Overall, nf-gwas is built on our knowledge of executing GWAS for many years, designed for thousands of phenotypes with scalable reporting features, is well tested and already attracted many other GWAS users.

## Conclusion

In the work at hand, we presented nf-gwas, a scalable pipeline based on Nextflow to run GWAS analysis in a distributed environment. The pipeline consists of several pre- and post-processing steps and currently uses REGENIE as a regression model for association analysis. The pipeline is built on our experience in the field of GWAS analysis from many years and helped our institute to focus more on data interpretation instead of workflow implementation details. We especially focused on high parallelization of data-intensive steps, whereas the overall parallelization level can be managed by the end-user enabling a maximum of flexibility. The pipeline is well-documented, outputs interactive web reports including millions of data points, provides tutorials for new users and is tested using the unit-style testing framework nf-test. Our HPC benchmarks and cost estimations on AWS show that the pipeline scales well and will simplify the executions of GWAS for the years to come.

## Acknowledgments

The authors would like to thank Adriana Koller and Azin Kheirkhah from the Institute of Genetic Epidemiology (Medical University of Innsbruck) for testing and their critical feedback. We also want to thank all GitHub contributors for improving the pipeline in various ways. A special thanks goes to the teams of Seqera Labs and nf-core for helping us to make the pipeline cloud-ready. We also appreciate the continuous support from the IT team of the Medical University of Innsbruck for providing the HPC cluster to run our large-scale experiments.

## Software Availability

nf-gwas is available at https://genepi.github.io/nf-gwas under the MIT license and requires Java 11 or higher for local execution.

## Author Contributions

LF and SS supervised the project and implemented the software. JSR, LF, SDM, and SS wrote the manuscript. CL, JSR, MF, MM and SDM tested the pipeline and contributed to the software. CF and FK provided extensive feedback to experiments, manuscript, and figures. All authors read and approved the final manuscript.

## Resources

nf-gwas Repository - https://github.com/genepi/nf-gwas

nf-gwas Documentation - https://genepi.github.io/nf-gwas

nf-test - https://github.com/askimed/nf-test

Pan-UK Biobank - https://pan.ukbb.broadinstitute.org

genomic-utils library - https://github.com/genepi/genomic-utils

GENCODE - https://www.gencodegenes.org

PheWeb - https://resources.pheweb.org

nf-core - https://nf-co.re

Nextflow Tower - https://tower.nf

Michigan Imputation Server - https://imputationserver.sph.umich.edu

GWAS catalog - https://www.ebi.ac.uk/gwas/home

