## Supplemental Material for "Performing highly parallelized and reproducible GWAS analysis on biobank-scale data"

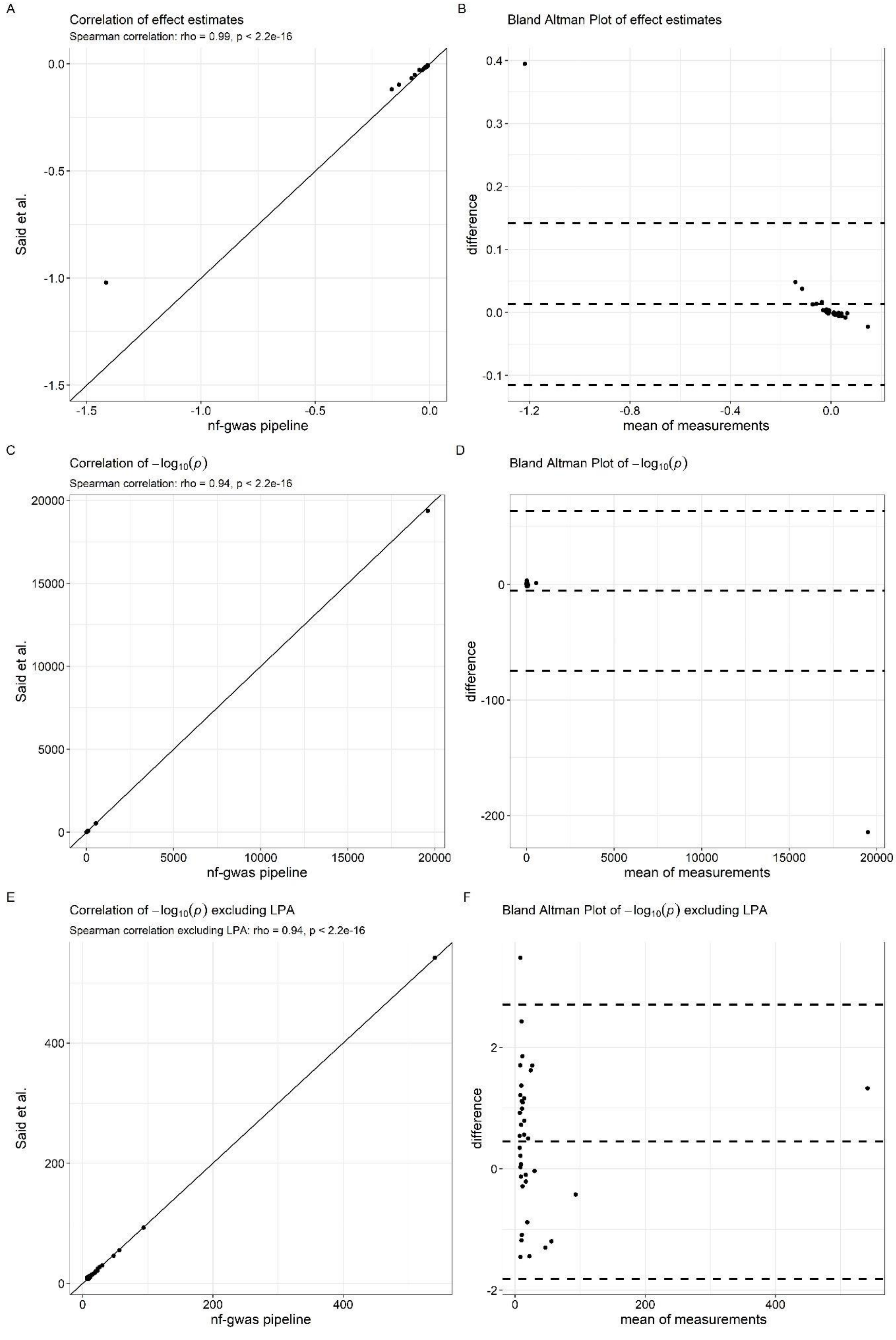
A
Correlation of effect estimates
Spearman correlation: rho = 0.99, p < 2.2e-16
Said et al.
nf-gwas pipeline
0.0
-0.5
-1.0
-1.5
B
Bland Altman Plot of effect estimates
difference
mean of measurements
0.4
0.3
0.2
0.1
0.0
-0.1
-1.2
-0.8
-0.4
0.0
C
Correlation of $-\log_{10}(p)$
Spearman correlation: rho = 0.94, p < 2.2e-16
Said et al.
nf-gwas pipeline
20000
15000
10000
5000
0
D
Bland Altman Plot of $-\log_{10}(p)$
difference
mean of measurements
0
-100
-200
0
5000
10000
15000
20000
E
Correlation of $-\log_{10}(p)$ excluding LPA
Spearman correlation excluding LPA: rho = 0.94, p < 2.2e-16
Said et al.
nf-gwas pipeline
400
200
0
F
Bland Altman Plot of $-\log_{10}(p)$ excluding LPA
difference
mean of measurements
2
0
-2
0
200
400

***Figure S1: Comparison of sentinel SNPs of the Lp(a) GWAS by Said et al. to our pipeline.***

*Plots A-D contain 37 variants (rs139970591 and rs531017143 were not available in our GWAS results) and plots E and F additionally exclude rs10455872 (LPA). A, C and E depict correlation plots and the black line denotes equal values between both studies. B, D and F depict Bland Altman plots depict mean and difference with x = Said et al. and y = nf-gwas pipeline of effect estimates on RINTed Lp(a) in B and $-log_{10}(p)$ in D and F and the dashed lines represent mean ± 2 SDs of differences (95% limits of agreement).*

***Table S1:*** *Comparison of effect estimates and -log*$_{10}$*(p) for tophits of the GWAS on RINTed Lp(a) between our pipeline and Said and colleagues. Chr: Chromosome; LOG10P : -log*$_{10}$*(p)*

| | | | nf-gwas | | | | | | | Said et al. | |
|---|---|---|---|---|---|---|---|---|---|---|---|
| **rsID** | **Chr** | **Gene** | **A0** | **A1** | **A1FREQ** | **INFO** | **BETA** | **SE** | **LOG10P** | **BETA** | **LOG10P** |
| rs182050989 | 1 | NUDC | T | C | 0.97 | 0.99 | 0.03 | 0.01 | 8.67 | 0.03 | 11.10 |
| rs11591147 | 1 | PCSK9 | T | G | 0.98 | 1.00 | 0.05 | 0.01 | 11.21 | 0.04 | 12.30 |
| rs599839 | 1 | CELSR2, PSRC1 | A | G | 0.24 | 1.00 | -0.02 | 0.00 | 23.38 | -0.02 | 25.00 |
| rs934197 | 2 | APOB | A | G | 0.67 | 1.00 | -0.02 | 0.00 | 19.80 | -0.01 | 20.30 |
| rs1047891 | 2 | CPS1 | A | C | 0.68 | 1.00 | 0.02 | 0.00 | 30.13 | 0.02 | 30.10 |
| rs115562858 | 3 | IP6K2, PRKAR2A | C | G | 0.98 | 0.98 | 0.04 | 0.01 | 8.27 | 0.04 | 8.30 |
| rs77601270 | 3 | RBM15B, VPRBP | C | G | 0.98 | 0.98 | 0.04 | 0.01 | 9.45 | 0.04 | 9.52 |
| rs645040 | 3 | MSL2 | T | G | 0.23 | 1.00 | 0.02 | 0.00 | 22.74 | 0.02 | 21.30 |
| rs2045592 | 4 | AFF1 | G | C | 0.62 | 1.00 | -0.01 | 0.00 | 8.18 | -0.01 | 8.40 |
| rs1908961 | 4 | ADH7 | G | A | 0.79 | 0.99 | -0.01 | 0.00 | 10.54 | -0.01 | 12.40 |
| rs114816312 | 4 | PLA2G12A | T | C | 0.99 | 1.00 | 0.07 | 0.01 | 9.88 | 0.06 | 11.00 |
| rs143500414 | 6 | TULP4 | C | T | 0.98 | 0.87 | -0.07 | 0.01 | 10.31 | -0.05 | 11.30 |
| rs141117870 | 6 | FNDC1 | T | C | 0.98 | 0.96 | -0.17 | 0.01 | 93.43 | -0.12 | 93.00 |
| rs10455872 | 6 | LPA | G | A | 0.93 | 1.00 | -1.42 | 0.00 | 19593.70 | -1.02 | 19379.40 |
| rs191369471 | 6 | PARK2 | G | C | 0.99 | 0.85 | -0.13 | 0.02 | 16.61 | -0.10 | 16.40 |
| rs34115226 | 7 | NFE2L3 | CG | C | 0.20 | 0.99 | 0.01 | 0.00 | 6.52 | 0.01 | 10.00 |
| rs35797675 | 7 | BAZ1B | G | T | 0.79 | 0.98 | 0.01 | 0.00 | 8.97 | 0.01 | 9.70 |
| rs4841132 | 8 | PPP1R3B | G | A | 0.09 | 1.00 | -0.02 | 0.00 | 11.09 | -0.02 | 10.00 |
| rs17411113 | 8 | LPL | G | C | 0.90 | 1.00 | -0.03 | 0.00 | 19.18 | -0.02 | 18.30 |
| rs61887548 | 11 | CHKA | C | T | 0.98 | 1.00 | 0.03 | 0.01 | 8.85 | 0.03 | 10.22 |
| rs2657878 | 12 | MIP, SPRYD4, GLS2 | T | C | 0.82 | 0.99 | 0.01 | 0.00 | 9.15 | 0.01 | 7.70 |
| rs139097404 | 15 | STRC, CATSPER2 | C | T | 0.98 | 0.97 | 0.06 | 0.01 | 25.70 | 0.05 | 27.40 |
| rs1532085 | 15 | ALDH1A2 | G | A | 0.39 | 1.00 | 0.01 | 0.00 | 7.34 | 0.01 | 9.05 |
| rs247616 | 16 | CETP | T | C | 0.68 | 1.00 | 0.03 | 0.00 | 47.30 | 0.02 | 46.00 |
| rs1110572 | 16 | PRMT7, SMPD3 | G | A | 0.88 | 1.00 | -0.02 | 0.00 | 11.81 | -0.02 | 11.52 |
| rs34042070 | 16 | TXNL4B, HP, HPR | G | C | 0.81 | 0.99 | -0.02 | 0.00 | 13.54 | -0.02 | 14.70 |
| rs139970591 | 17 | PEMT | NA | NA | NA | NA | NA | NA | NA | 0.01 | 12.00 |
| rs3785549 | 17 | PSMD3 | C | T | 0.46 | 0.99 | -0.01 | 0.00 | 7.08 | -0.01 | 8.00 |
| rs1801689 | 17 | APOH | C | A | 0.97 | 1.00 | -0.08 | 0.01 | 56.49 | -0.07 | 55.30 |
| rs77542162 | 17 | ABCA6 | G | A | 0.98 | 1.00 | -0.05 | 0.01 | 13.96 | -0.03 | 14.52 |
| rs2292642 | 17 | PGS1 | T | C | 0.40 | 1.00 | -0.01 | 0.00 | 6.98 | -0.01 | 7.52 |
| rs874492 | 19 | CLEC4M | T | A | 0.68 | 0.99 | -0.01 | 0.00 | 7.05 | -0.01 | 7.40 |
| rs138294113 | 19 | LDLR | T | C | 0.88 | 1.00 | 0.02 | 0.00 | 16.62 | 0.02 | 16.52 |
| rs1065853 | 19 | TOMM40, APOE, APOC1 | T | G | 0.92 | 1.00 | 0.16 | 0.00 | 540.67 | 0.13 | 542.00 |
| rs35866622 | 19 | FUT2, MAMSTR, RASIP1 | T | C | 0.52 | 0.99 | -0.01 | 0.00 | 13.91 | -0.01 | 14.70 |
| rs8736 | 19 | TMC4, MBOAT7 | T | C | 0.56 | 0.99 | -0.01 | 0.00 | 10.33 | -0.01 | 9.15 |
| rs73075609 | 20 | GPCPD1 | T | C | 0.97 | 0.98 | -0.03 | 0.01 | 9.28 | -0.03 | 9.15 |
| rs531017143 | 20 | ZGPAT, LIME1, SLC2A4RG | NA | NA | NA | NA | NA | NA | NA | -0.01 | 10.40 |

***Table S2:*** *Comparison of features of existing pipelines.*

| | nf-gwas | nf-gwas-pipeline | H3AGWAS | BIGwas |
|---|---|---|---|---|
| **GitHub** | https://github.com/genepi/nf-gwas | https://github.com/montilab/nf-gwas-pipeline | https://github.com/h3abionet/h3agwas | https://github.com/ikmb/gwas-assoc |
| **Association software** | REGENIE | GENESIS, GMMAT | gcta, plink, gemma, Bolt-LMM, REGENIE, saige, fast-lmm | PLINK, SAIGE |
| **Input formats** | VCF, BGEN | VCF or gds | depends on software | VCF |
| **Quantitative traits** | yes | yes | yes | yes |
| **Binary traits** | yes | yes | yes | yes |
| **>1 phenotype** | yes | no | yes | no |
| **Step 1 QC** | yes | not applicable | not applicable/no | Not applicable |
| **Single-variant testing** | yes | yes | yes | yes |
| **Rare-variant/gene-based testing** | yes | yes | no | no |
| **GxE testing** | yes | no | yes | no |
| **GxG testing** | yes | no | no | no |
| **Conditional analysis** | yes | no | no | no |
| **longitudinal analysis** | no | yes | no | no |
| **Report** | yes | yes | yes | yes |
| **Interactive report** | yes | no | no | no |
| **Tabix-indexed (LocusZoom ready)** | yes | no | no | no |
| **Annotations** | nearest gene + rsID | ANNOVAR | In separate pipeline | no |
| **Lift over** | yes | no | no | In separate QC pipeline |

| | | | | |
|---|---|---|---|---|
| **option** | | | | |
| **PC generation** | no | yes | no | yes |
| **Addition of own PCs** | yes | yes | yes | no |
| **Software tests** | yes (nf-test) | no | no | no |
